# Altered lipid metabolism marks glioblastoma stem and non-stem cells in separate tumor niches

**DOI:** 10.1101/2020.09.20.304964

**Authors:** Sajina Shakya, Anthony D. Gromovsky, James S. Hale, Arnon M. Knudsen, Briana Prager, Lisa C. Wallace, Luiz O. F. Penalva, Pavlina Ivanova, H. Alex Brown, Bjarne W. Kristensen, Jeremy N. Rich, Justin D. Lathia, J. Mark Brown, Christopher G. Hubert

**Affiliations:** Department of Biomedical Engineering, Cleveland Clinic Lerner Research Institute, Cleveland OH, USA; Cardiovascular and Metabolic Sciences, Cleveland Clinic Lerner Research Institute, Cleveland OH, USA; Case Comprehensive Cancer Center, Case Western Reserve University, Cleveland OH, USA; Department of Pathology, Odense University Hospital, Odense, Denmark; Department of Clinical Research, University of Southern Denmark, Odense, Denmark; Cleveland Clinic Lerner College of Medicine, Cleveland Clinic, Cleveland OH, USA; Medical Scientist Training Program, Case Western Reserve School of Medicine, Cleveland OH, USA; Children’s Cancer Research Institute, University of Texas Health Sciences Center San Antonio, San Antonio, TX, USA; Department of Pharmacology, Vanderbilt University School of Medicine, Nashville, Tennessee; Department of Medicine, University of California San Diego, San Diego CA, USA

**Keywords:** Glioblastoma, organoid, tumor heterogeneity, lipid droplets, cancer stem cell

## Abstract

**Background:** Glioblastoma (GBM) is marked by cellular heterogeneity, including metabolic heterogeneity, that varies among cellular microenvironments in the same tumor. Altered cellular metabolism in cancer is well-established, but how lipid metabolism is altered to suit different microenvironmental conditions and cellular states within a tumor remains unexplored.

**Methods:** We assessed GBM organoid models that mimic the transition zone between nutrient-rich and nutrient-poor pseudopalisading/perinecrotic tumor zones and performed spatial RNA-sequencing of cells to interrogate lipid metabolism. Using targeted lipidomic analysis, we assessed differences in acutely enriched cancer stem cells (CSCs) and non-CSCs from multiple patient-derived models to explore the link between the stem cell state and lipid metabolism.

**Results:** Spatial analysis revealed a striking difference in lipid content between microenvironments, with lipid enrichment in the hypoxic organoid cores and the perinecrotic and pseudopalisading regions of primary patient tumors. This was accompanied by regionally restricted upregulation of hypoxia-inducible lipid droplet-associated (HILPDA) gene expression in organoid cores and in clinical GBM specimens, but not lower-grade brain tumors, that was specifically localized to pseudopalisading regions of patient tumors. CSCs have low lipid droplet accumulation compared to non-CSCs in organoid models and xenograft tumors, and prospectively sorted lipid-low GBM cells are functionally enriched for stem cell activity. Targeted lipidomic analysis revealed that CSCs had decreased levels of major classes of neutral lipids compared to non-CSCs but had significantly increased polyunsaturated fatty acid production due to high fatty acid desaturase (FADS1/2) expression.

**Conclusions:** Our data demonstrate that lipid metabolism is differentially altered across GBM microenvironments and cellular hierarchies, providing guidance for targeting of these altered lipid metabolic pathways.

**Key points:**

1. GBM cells in nutrient-poor tumor regions have increased accumulation of lipid droplets.
2. CSCs have reduced lipid content compared to non-CSCs.
3. GBM CSCs and non-CSCs have disparate lipid metabolisms that may be uniquely targetable.

**Importance of the Study:** Metabolic targeting has long been advocated as a therapy against many tumors including GBM, and it remains an outstanding question whether cancer stem cells (CSCs) have altered lipid metabolism. We demonstrated striking differences in lipid metabolism between diverse cell populations from the same patient. These spatially and phenotypically distinct lipid phenotypes occur clinically in the majority of patients and can be recapitulated in laboratory models. Lipidomic analysis of multiple patient-derived models shows a significant shift in lipid metabolism between GBM CSCs and non-CSCs, suggesting that lipid levels may not be simply a product of the microenvironment but also may be a reflection of cellular state. Our results suggest that therapeutic targeting of GBM lipid metabolism must consider multiple separate tumor cell populations to be effective, and we provide a methodologic framework for studying these metabolically diverse cellular populations.

## INTRODUCTION

Glioblastoma (GBM) is the most common primary malignant brain tumor in adults. Despite aggressive standard treatment strategies including surgical resection followed by radiation and chemotherapy, the median survival for patients with GBM is approximately 15 months from the time of diagnosis.^1^ A key challenge to GBM treatment is the intratumoral heterogeneity at both the cellular and microenvironmental levels.^2,3^ Maintenance of heterogeneity may be driven by a population of cells within the tumor termed cancer stem cells (CSCs), which are highly plastic and responsive to their environment and hold self-renewal and tumor initiation capacity.^4,5^ Critically for treatment, this diversity within the cell population means that while cells from one microenvironment or cellular state may respond to a therapy, others may not, resulting in therapeutic resistance of the overall tumor. An effective GBM treatment plan should therefore require combination of several approaches that could target distinct aspects of tumor cells in these microenvironments.

In cancers, cells adapt through a metabolic shift to sustain the higher energy demand of oncogenic growth, to conserve self-renewing CSCs, and to survive under unfavorable microenvironmental conditions such as hypoxia.^6^ Tumor cells often undergo metabolic reprogramming, also called the “Warburg Effect”, where they primarily use glycolysis to produce cellular ATP even in high-oxygen environments. This is in contrast to normal non-proliferating cells, which typically generate 90% of their ATP in the mitochondria. The altered metabolism of tumor cells is especially efficient in the hypoxic microenvironment of GBM tumors, where cells do not have access to the resources necessary to generate ATP via normal oxidative phosphorylation^7,8^. This results in differential regulation of various metabolic genes that function as part of the glucose, lipid, glutamine, and nucleotide metabolic pathways.^9^ Recently, lipid metabolism has emerged as a potential therapeutic target to treat gliomas, including GBM^10,11^, and brain metastases^12^. Lipids are a group of fat-soluble organic compounds that act as cell membrane structural components, energy storage units, and signaling molecules.^13^ Common lipid types include fatty acids, phospholipids and cholesterol. Lipid metabolism is abnormally regulated in gliomas compared to normal cells, with changes in the expression of lipid-related genes such as SREBP1 and FAS, which results in altered lipid composition and lipogenesis to keep up with energy demands.^10,14,15^

Lipid droplets are cytosolic organelles that, among other functions, serve as a storage medium for the fatty acids used as an energy source to maintain rapid proliferation in unfavorable microenvironments. Lipid droplet formation particularly occurs under stressful conditions such as hypoxia and nutrient deprivation.^16^ Accumulation of lipid droplets has been observed in a variety of cancers, including hepatic cancer, lung cancer, breast cancer, and gliomas, and is an important regulator of critical facets of cancer including angiogenesis, inflammatory responses, apoptosis and cell death, and hypoxia-mediated alterations of lipid metabolism.^17^ Although lipid droplet accumulation has been observed in hypoxic cells within several tumor types^18,19^, differential lipid metabolism between heterogeneous GBM cell types and microenvironments has remained unexplored. In this study, we analyzed spatial alterations in gene expression and lipid content to determine the metabolic alterations present in GBM CSCs.

## METHODS

### Human cell and organoid culture

Glioblastoma samples were obtained either directly from patients undergoing resection following written informed consent in accordance with protocol #2559 approved by the Cleveland Clinic Institutional Review Board or from collaborators as previously established patient-derived tumorsphere cultures. Patient tissue samples were either finely minced prior to organoid formation or were dissociated into single-cell suspensions, red blood cells were then removed by brief hypotonic lysis, and cells were counted for number and viability using trypan blue. Cells were cultured as tumorspheres in Neurobasal medium supplemented with 10ng/mL EGF (R&D Systems, Minneapolis MN), 10ng/mL bFGF (R&D Systems), B27 (Invitrogen, Carlsbad CA), glutamine (CCF media core), sodium pyruvate (Invitrogen), and antibiotics (Antibiotic-Antimycotic, Invitrogen) (“NBM complete”). All cells used in this work were patient-derived primary cultures, and all specimens were verified by comparison of short tandem repeat (STR) analysis performed periodically during the course of experimentation. Tumorspheres were used to form xenografts and harvested for analysis as previously described.^20^ For prospective stem cell sorting, xenografts were minced and digested with papain (Worthington) as previously described^21^, and dissociated cells were allowed to recover overnight prior to use. Following overnight recovery from papain digestion, dissociated xenograft GBM cells were magnetically sorted based on CD133 expression using magnetic beads (CD133/2 beads, Miltenyi). This approach has previously been validated to show differences in tumorigenic potential between CD133-positive and CD133-negative fractions.^20–24^ Organoids were formed as previously described^25^ by suspending tumor cells in 80% Matrigel (BD Biosciences, San Jose, CA) and forming 20 μl pearls on parafilm molds prior to culture. Organoids were cultured in 6-well or 10-cm plates with shaking in NBM complete media.

### Regional isolation of GBM organoids

Regional labeling and subsequent cell isolation from organoid layers was achieved using a 20 μM final concentration of CellTracker Blue CMAC Dye (Invitrogen, #C2110) in media. Mature organoids were incubated with dye for 2 hours at 37°C with shaking to allow outer layer labeling. Desired labeling depth was verified using a confocal microscope and a compatible imaging dish (MatTek #P35GC-1.0-14-C). After labeling, organoids were finely chopped and dissociated using Accutase (FisherSci, #ICN1000449) at 4°C for 15 minutes and then heated to 37°C for another 10 minutes. Cells were then single-cell filtered, and live cells were isolated by fluorescence-activated cell sorting (FACS) using 1 μM Calcein-AM (Invitrogen, #C3099MP) and 1:2000 TO-PRO3 (Invitrogen, #T3605) according to the manufacturer’s protocols. Cell sorting and analysis were performed using a BD FACS ARIA II flow cytometer.

### RNA-sequencing analysis

Total RNA was extracted using TRIzol reagent (Life Technologies) and purified with phenol-chloroform extraction including the use of PhaseLock tubes (5PRIME). Samples were prepared for RNA-seq according to the manufacturer’s instructions (Illumina) and sequenced using 101-bp paired-end chemistry on a HiSeq-2000 machine in the UTHSCSA Genomic Facility. FASTQ files were trimmed with TrimGalore, which implements Cutadapt^26^ and FastQC^27^, to remove low quality reads and trim adapters. Reads were aligned to Gencode v29 using Salmon^28^ with correction for GC bias, positional bias and sequence-specific bias. The R/Bioconductor package tximport^29^ was used to generate TPM values. Low-expressed genes without at least 1 count in 4 samples were excluded from analysis. Comparisons were performed using DESeq2^30^ on raw counts. PCA plots were generated using the ‘prcomp’ function from the R/Bioconductor package ‘stats’ with default arguments. GSEA analysis was performed using mSigDB (Broad Institute).

### Limiting dilution assay

To determine tumorsphere-propagating potential, live cells were sorted into wells of 96-well plates at concentrations ranging from 1 to 32 cells per well. Cells were then grown in a 37°C tissue culture incubator in culture media for 14 days. The presence or absence of spheres in each well was then assessed and analyzed using the ELDA analysis tool (http://bioinf.wehi.edu.au/software/elda/) to calculate stem cell frequencies.^31^

### Oil Red O histochemistry

For organoid Oil Red O staining, whole organoids were fixed in 4% paraformaldehyde, cryoprotected in 30% sucrose, and snap frozen in OCT using an isopentane bath chilled with dry ice. Tissue sections with a thickness of 10 μm were cut on a cryomicrotome and mounted on glass slides. Oil Red O staining was performed by the Cleveland Clinic Lerner Research Institute imaging core following standard core protocols using commercial control slides. Slides were digitized with a Leica Aperio digital slide scanner (Leica). For primary patient GBM samples, freshly resected GBM tissue specimens from the Department of Neurosurgery, Odense University Hospital, Odense, Denmark, were frozen at −40°C using the MCC cryoembedding compound and PrestoCHILL device (Milestone). Tissue sections with a thickness of 8 μm were cut on a cryomicrotome and mounted on glass slides. The slides were left to dry for 15 minutes at room temperature and fixed for one hour. Following fixation, slides were washed three times with deionized water and incubated with Oil Red O staining solution (Fluka, CI26125, dissolved in 60% triethyl phosphate) for 30 minutes. After staining, slides were washed with deionized water three times and counterstained with Mayer’s hematoxylin (Merck). Slides were then rinsed with deionized water for 5 minutes and mounted with a coverslip using Aquatex mounting medium. Finally, slides were digitalized with the NanoZoomer 2.0HT digital image scanner (Hamamatsu, Japan). The use of tissue specimens was approved by the Danish Data Inspection Authority (approval number 16/11065) and the Regional Scientific Ethical Committee of the Region of Southern Denmark (approval number S-20150148).

### Targeted Lipidomic Profiling

Quantification of neutral lipids and glycerophospholipids was conducted as previously described.^32,33^ Briefly, approximately 10 mg of frozen mouse liver was homogenized in 800 mL ice-cold 0.1 N HCl:CH_3_OH (1:1) using a tight-fit glass homogenizer (Kimble/Kontes Glass, Vineland, NJ) for ~1 min on ice. The suspension was then transferred to cold 1.5 mL microfuge tubes (Laboratory Product Sales, Rochester, NY) and vortexed with 400 mL cold CHCl_3_ for 1 min. The extraction proceeded with centrifugation (5 min, 4°C, 18,000g) to separate the two phases. The lower organic layer was collected, and the solvent was evaporated. The resulting lipid film was dissolved in 100 mL isopropanol:hexane:100 mmol/L NH_4_CO_2_H(aq) (58:40:2) (mobile phase A). Quantification of glycerophospholipids was achieved by the use of a liquid chromatography–mass spectrometry technique using synthetic (non–naturally occurring) diacyl and lysophospholipid standards. Typically, 200 ng of each odd-carbon standard was added per 10–20 mg tissue. Glycerophospholipids were analyzed on an Applied Biosystems/MDS SCIEX 4000 Q TRAP hybrid triple quadrupole/linear ion trap mass spectrometer (Applied Biosystems, Foster City, CA) and a Shimadzu high-pressure liquid chromatography system with a Phenomenex Luna Silica column (2 3 250 mm, 5 mm particle size) using a gradient elution. The identification of the individual species, achieved by liquid chromatography-tandem mass spectrometry, was based on their chromatographic and mass spectral characteristics. This analysis allows identification of the two fatty acid moieties but does not determine their position on the glycerol backbone (sn-1 vs. sn-2). Triacylglycerol (TAG), diacylglycerol (DAG), and monoacylglycerol (MAG) from frozen mouse liver tissue (10–15 mg) were extracted by homogenizing tissue in the presence of internal standards (500 ng each of 14:0 MAG, 24:0 DAG, and 42:0 TAG) in 2 mL PBS and extracting with 2 mL ethyl acetate: trimethylpentane (25:75). After drying the extracts, the lipid film was dissolved in 1 mL hexane:isopropanol (4:1) and passed through a bed of 60 Å Silica gel to remove the remaining polar phospholipids. Solvent from the collected fractions was evaporated, and lipid film was redissolved in 100 mL CH_3_OH:CHCl_3_ (9:1) containing 10 mL of 100 mmol/L CH_3_COONa for mass spectrometry analysis as described previously.^32,33^

## RESULTS

### GBM organoids mimic the pathologic transition zones and molecular heterogeneity of GBM patient tumors

GBM tumors have a complex microenvironment defined by histologic hallmarks of angiogenesis and pseudopalisading necrosis, and these two anatomic features harbor distinct GBM cell populations. CSCs are particularly enriched in the perivascular niche of glioblastoma tumors.^34^ Current culture models fail to replicate the complex microenvironments of CSCs, limiting our ability to study and therapeutically target GBM. We previously developed 3D patient-derived GBM organoids that show a heterogeneous mix of cells forming zones comparative to the pathologic transition zones in patient tumors.^35^ GBM organoids can be conceptually divided into two zones - an outer cell-dense rim consists of dividing cells in a high-oxygen and high-nutrient environment provided by the nearby media, mimicking the conditions of the tumor perivascular niche, and a hypoxic core with necrotic cells relatively deprived of the nutrient media components, which phenotypically mimics the hypoxic and perinecrotic regions of GBM tumors (Fig. 1A).

**Figure 1.**
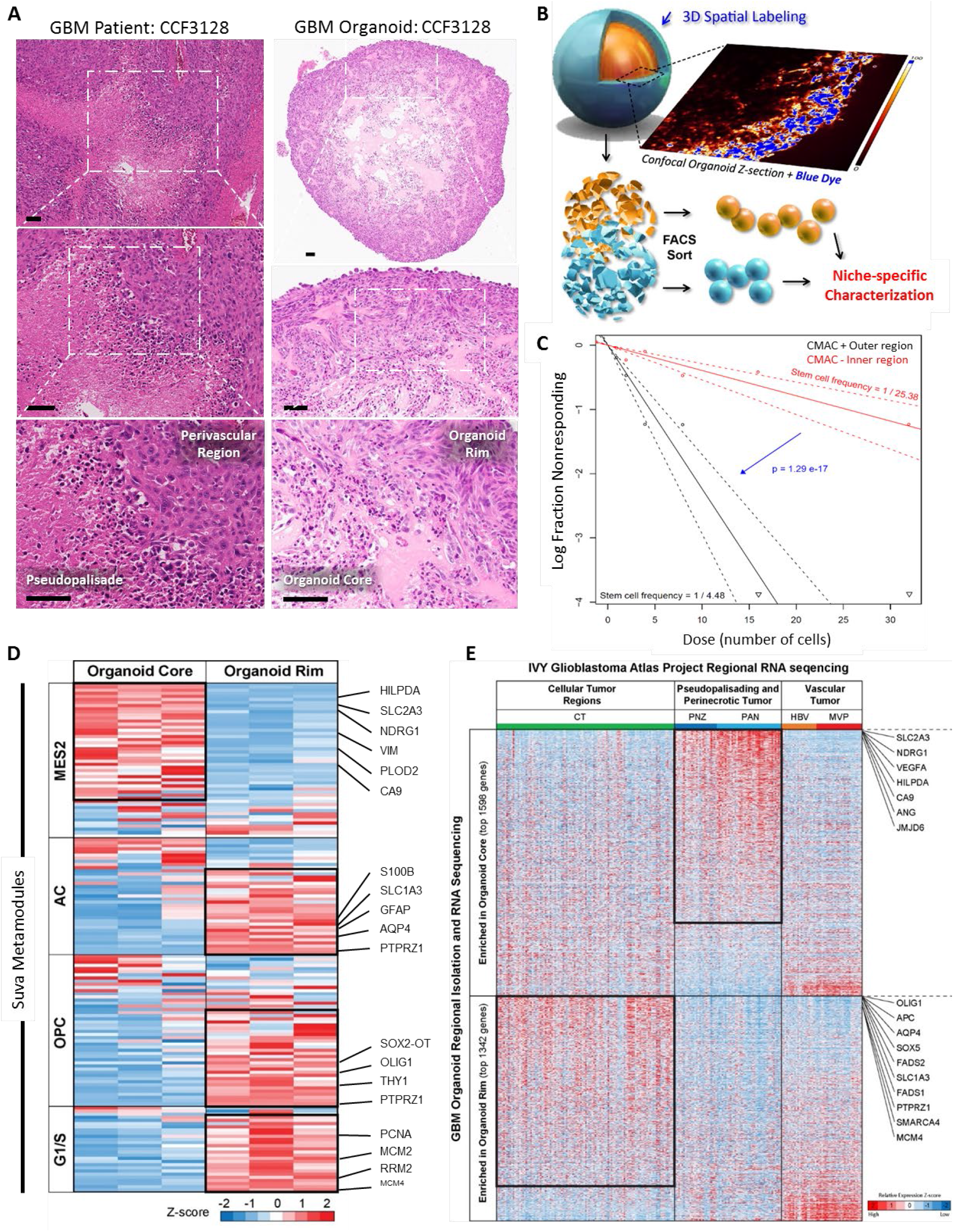
GBM organoids mimic the pathologic transition zones and molecular heterogeneity of GBM patient tumors. (A) H&E staining of GBM 3D organoids (right panel) reveals histological zones comparable to GBM primary patient tumors (left panel). The perivascular region and hypoxic core in primary patient tumors (left panel) are mimicked by the organoid proliferative rim and hypoxic core regions (right panel), respectively. (B) To compare the molecular signature of these histological regions, the organoids were stained whole to label the entire outer rim region, and single cells were isolated. (C) Limiting-dilution assays showed that the organoid proliferative rim is functionally enriched for stem cells compared to the hypoxic core. Calculated stem cell frequencies and 95% confidence intervals are shown. (D) Upon RNA-seq analysis of single cells, distinct cell-type signatures were found to be enriched within spatially separate niches in the organoids. (E) Mapping expression in organoids to the regional Ivy GAP database showed region-specific enrichment. *Scale bar = 100 μm*

To determine whether these ex vivo culture regions molecularly mimic the corresponding patient GBM tumor regions, we developed a method to three-dimensionally label and sort live GBM cells from our organoid cultures (Fig. 1B). Whole organoids were incubated with a blue lipophilic dye for an optimized period that we empirically determined to be sufficient to label only the outer proliferative niche as viewed by live confocal z-section imaging. Organoids were then dissociated and sorted via FACS to separate these spatially distinct cell populations. Cells in the outer rim display enhanced self-renewal, a CSC hallmark, compared to cells within the core, as determined by limiting-dilution assay (Fig. 1C).

We performed spatially defined RNA-sequencing of the different regions of GBM organoids to investigate the gene expression in each niche population. Recently, Neftel et. al defined single-cell heterogeneity of patient GBM cell populations as being dominated by clusters of genes that change in relationship to each other, called meta-modules.^36^ We compared these gene expression signatures to these meta-module gene signatures from single cell RNA sequencing (scRNA-seq) of clinical GBM. We found distinct cell-type signatures enriched within spatially separate niches in our organoids (Fig. 1D). As anticipated, the cells from the organoid core were enriched for hypoxia hallmark genes as determined by gene set enrichment analysis (GSEA; FDR q value = 3.22 x 10^-16^) and displayed a signature of the hypoxia-dependent mesenchymal-like 2 (MES2) meta-module defined by scRNA-seq of patient tumor cells. In contrast, cells from the organoid rim were enriched for genes found in the astrocyte-like (AC) and oligodendrocyte precursor cell (OPC)-like meta-modules (Fig. 1D). Additionally, the cells from the organoid rim highly expressed genes related to the G1/S meta-module, indicating high proliferation in this niche. Overall, the expression profiles within our spatially segregated organoid microenvironments represent 3 out of 4 categories of discrete GBM cell types identified in clinical tumors by Neftel et al.^36^

While these data provide resolution to connect the diverse cell types present in GBM organoids with individual cell signatures from clinical tumors, the single-cell sequencing data cannot be directly traced to spatial tumor subregions. To correlate the niche-specific gene expression in organoids and the gene expression of different regions of primary GBM tumors, we reflected our sequencing data upon subregion sequencing data from 41 patient samples in the Ivy Glioblastoma Atlas Project (Ivy GAP) database^37^ (Fig. 1E). We found that genes significantly enriched in the organoid core were specifically highly expressed in the pseudopalisading and perinecrotic tumor regions of primary GBM. Conversely, the organoid rim was found to be enriched for genes expressed in the cellular tumor regions and depleted of those expressed in regions of hypoxia. Taken together, these data show that the 3D environments within GBM organoids recapitulate at least part of the cellular and microenvironmental diversity within primary GBM tumors at both a histologic and molecular level.

Genes that are differentially overexpressed in tumor compared to normal brain tissue may suggest a possible therapeutic window for a candidate therapeutic target in brain tumors. To determine genes from our analysis that may be upregulated in tumor compared to normal brain, and upregulated in increasing grades of tumor, we compared the TCGA, Gravendeel, Bao and Ivy GAP datasets using the GlioVis portal^38^. First, to validate our findings, we chose to look at differential expression of specific genes that represent the different regions, particularly highlighting representative genes from the three different meta-modules, including a well-established hypoxia marker, CAIX (Supplemental Fig. S1). We found that carbonic anhydrase 9 (CAIX) and vimentin (VIM) (Mes2 meta-module) were highly expressed in GBM brain tumor compared to non-tumor tissue, due in part to their increase pseudopalisading regions. This was reflected in GBM organoids where expression of the CAIX and VIM genes was likewise increased in the core region compared to the rim. Although the expression of the OLIG1 (OPC meta-module) and AQP4 (AC meta-module) genes was not significantly increased in GBM compared to non-tumor tissue, these genes are differentially expressed within GBM patient tumor regions and these differences are again recapitulated in the corresponding organoid regions. Taken together, the above findings demonstrate that we can recapitulate tumor-relevant cellular heterogeneity and maintain microenvironmentally regulated GBM cell behavior ex vivo.

### Lipid droplets accumulate in the perinecrotic and pseudopalisading zones of GBM patient tumors and the corresponding zones of GBM organoids

In addition to the above genes, we identified the gene hypoxia-inducible lipid droplet-associated (HILPDA), which encodes a protein necessary for lipid trafficking in cytosolic lipid droplets, to be consistently differentially expressed in the different regions of 3D GBM organoids and GBM clinical datasets (Fig 2A). HILPDA expression was significantly higher in brain tumor tissue compared to normal brain and higher in GBM compared to lower-grade brain tumors. Moreover, in primary GBM, HILPDA expression was significantly higher in the pseudopalisading region of the tumor. The increase in HILPDA expression in only GBM and not lower-grade brain tumors, combined with the specific increase in pseudopalisading regions, is consistent with brain tumor pathology as pseudopalisading necrosis is a defining diagnostic feature of GBM. We observed similar differences in the corresponding regions of GBM organoids: HILPDA expression was significantly higher in the core region of organoids compared to the rim (Fig 2A).

**Figure 2.**
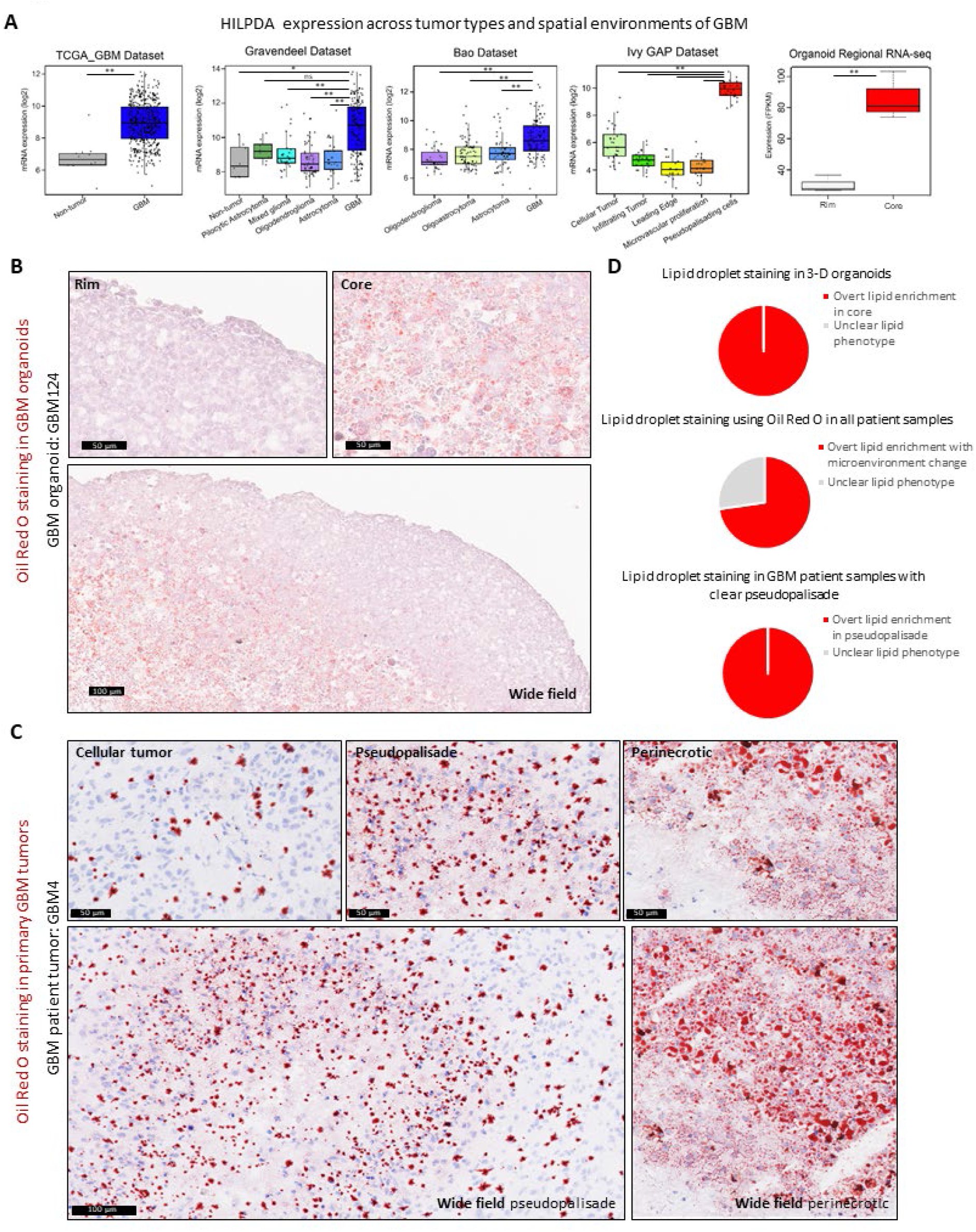
Lipid droplets accumulate in the perinecrotic and pseudopalisading zones of GBM patient tumors and the corresponding zones in organoids. (A) Publicly available databases show that HILPDA is consistently increased in GBM tumors and specifically enriched in hypoxic pseudopalisading cells, which is recapitulated by the organoid core. *\* p < 0.01; ** p < 0.001; ns, p > 0.05*. Lipid droplet staining with Oil Red O shows higher staining in the (B) organoid core and (C) pseudopalisading and perinecrotic regions of primary tumors. *Scale bar for wide field images = 100 μm and 50 μm for other images*. (D) Pie charts representing the findings in 11 patient tumors and 9 organoids.

The increase in HILPDA expression, combined with an increase in hallmark adipogenesis genes (p = 2.45 x 10^-5^, FDR q-value = 1.36 x 10^-4^) and cholesterol homeostasis genes (p = 2.16 x 10^-5^, FDR q-value = 1.36 x 10^-4^) in the organoid core compared to the rim, encouraged us to investigate whether there was any lipid accumulation phenotype that would indicate an overall alteration in lipid metabolism and storage. We therefore stained sections from primary GBM samples and lab-grown GBM organoids with a lysochrome diazo dye, Oil Red O, for histological visualization of lipid droplets. We observed notable differences in lipid droplet staining within the two regions of organoids. Oil Red O staining was concentrated in the cells of the core region of GBM organoids, indicating accumulation of lipid droplets in these cells, whereas the cells in the rim region were devoid of the stain (Fig. 2B, Supplemental Fig. S2). To determine whether this ex vivo phenotype is representative of human tumors, we further analyzed multiple primary patient GBM sections. Similar to what was observed in organoids, the cells in the corresponding pseudopalisading and perinecrotic regions of primary tumors specifically stained for the Oil Red O dye, while the cells in the cellular tumor region lacked the stain (Fig. 2C, Supplemental Fig. S3). This staining pattern held true for the vast majority (73%) of samples (Fig. 2D). These results demonstrate a generalizable phenomenon of altered lipid metabolism and storage between different regions of GBM tumor.

### Differential lipid accumulation marks GBM CSC and non-CSC populations

As the rim region of GBM organoids is enriched for CSCs and the hypoxic core has limited CSCs (Fig. 1C), we investigated whether lipid droplet accumulation is associated with stem cell phenotype. We chose to utilize *in vivo* patient-derived xenograft (PDX) models for this purpose as these represent perhaps the most realistic and cellularly diverse recapitulation of human tumor microenvironments. We collected and dissociated GBM tumor xenografts to obtain a heterogeneous mix of single tumor cells. We then magnetically sorted the cells into populations of CD133-positive cancer stem cells (CSCs) and CD133-negative non-stem cancer cells (non-CSCs) (Fig. 3A). Upon flow cytometric analysis, we observed increased fluorescence in the non-CSC populations of all three PDX specimens for both lipid dyes tested (Fig. 3C). Oil Red O dye staining of fixed sorted cells also confirmed this result (Fig. 3B).

**Figure 3.**
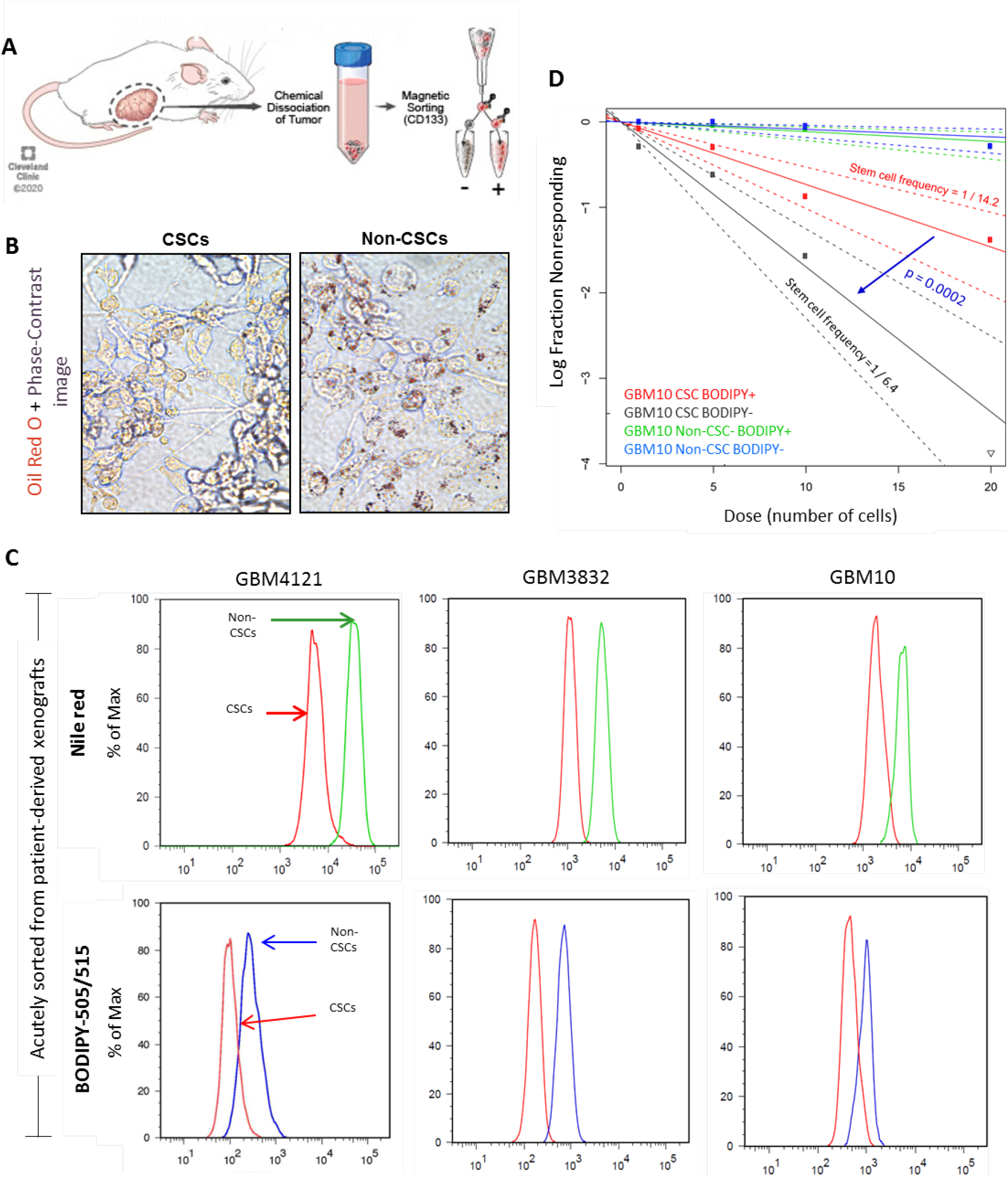
GBM CSCs and non-CSCs have differential levels of lipid accumulation. (A) Dissociated cells from GBM PDX models were stained for the stem cell marker CD133 and magnetically sorted. (B) Oil Red O staining was higher in the cultured non-CSC population than in the CSC population. (C) The CD133-negative non-CSC population showed increased fluorescence compared to CD133-positive CSCs for both Nile red and BODIPY lipid-specific fluorescent dyes. (D) Magnetically sorted CSC and non-CSC populations were subsequently FACS sorted into lipid-high and lipid-low populations based on BODIPY fluorescence (top and bottom 20%). Sphere-forming capability was determined by limiting-dilution assay. Non-CSCs showed uniformly low sphere-forming capability. BODIPY-low CSCs had significantly higher self-renewal capacity than their BODIPY-high counterparts (p=0.002). Calculated stem cell frequencies and 95% confidence intervals are shown.

To further validate this observation, we asked whether lipid content can enrich for CSCs with increased sphere-forming capability. We sorted dissociated GBM cells for CD133 as above, stained the CSCs and non-CSCs with BODIPY lipid dye, FACS sorted BODIPY-high and BODIPY-low populations, and tested their sphere-forming capability by limiting-dilution assay. Non-CSCs had uniformly low sphere-forming capability, as expected. However, BODIPY-low CSCs were functionally enriched for sphere-forming behavior compared to CSCs with high lipid content (Fig. 3D). We therefore conclude that CSCs are enriched among cells with reduced lipid droplet accumulation and that lipid accumulation may be an indicator of the CSC/non-CSC cell state.

### Lipidomic profiling of CSCs and non-CSCs from patient-derived samples reveals increased neutral lipid species in non-CSC populations

Non-CSCs have higher lipid accumulation in comparison to CSCs, but we cannot resolve individual lipid species by dye. We therefore investigated the differences in lipid metabolism between CSCs and non-CSCs using targeted lipidomic approaches. We isolated cells from 5 patient-derived PDX models, sorted for CD133+/- cell populations, and analyzed these as pools of CD133+ CSCs and CD133-non-CSCs (Fig. 4A) via targeted lipidomic profiling as previously described.^32,33^ We found that the high lipid content in the non-CSC population is due to a broad increase in lipids known to be preferentially stored in cytosolic lipid droplets. Neutral lipid species including diacylglycerol (DAG) and triacylglycerol (TAG) were significantly enriched in the non-CSCs (Fig. 4A, B). These data are consistent with the marked accumulation of cytosolic lipid droplets in non-CSCs.

**Figure 4.**
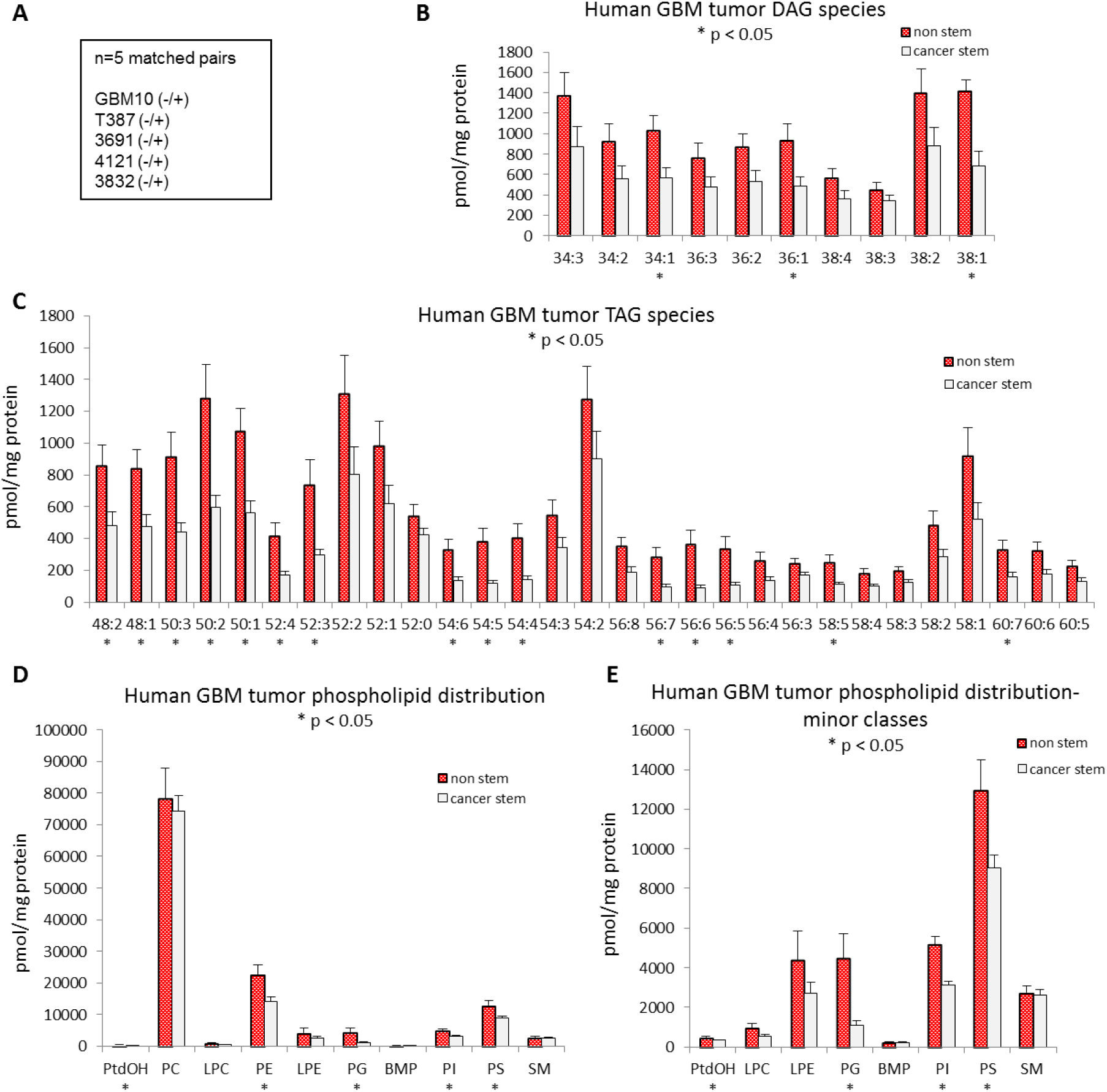
Lipid profiling of GBM stem and non-stem cells show differences in global lipid profiling. (A) Matched pairs (n=5) of magnetically sorted CD133-positive CSCs and CD133-negative non-CSCs were pooled and analyzed together for different classes of lipids using mass spectroscopy. GBM CSCs have decreased levels of the neutral lipid species (B) DAG and (C) TAG. Specific classes of phospholipids are altered in CSCs vs non-CSCs: (D) Overall phospholipid distribution and (E) Distribution of minor phospholipid classes. *\*p < 0.05*.

### GBM CSCs from patient-derived samples exhibit species-specific alterations in glycerophospholipids

Most, but not all (Fig. S4), molecular species of neutral lipids (DAGs and TAGs) were enriched in non-CSC populations. However, we observed numerous species-specific alterations in glycerophospholipid levels (Figs. 4–6). Analysis of total levels of glycerophospholipids revealed that CSCs exhibit modest decreases in minor phospholipid classes including phosphatidic acid (PA), phosphatidylethanolamine (PE), phosphatidylglycerol (PG), phosphatidylinositol (PI) and phosphatidyl (Fig. 4C, D). However, the most abundant class of glycerophospholipids, phosphatidylcholines (PC), was unaltered (Fig. 4C). When we examined molecular species within each class, there was a general decrease in the levels of longer-chain polyunsaturated fatty acid (PUFA) species of PS (Fig. 5A), PG (Fig. 5B), PE (Fig. 5D), and PC (Fig. 5E) in CSC populations, indicating that the esterification of PUFAs into these complex lipids may be selectively impaired.

**Figure 5.**
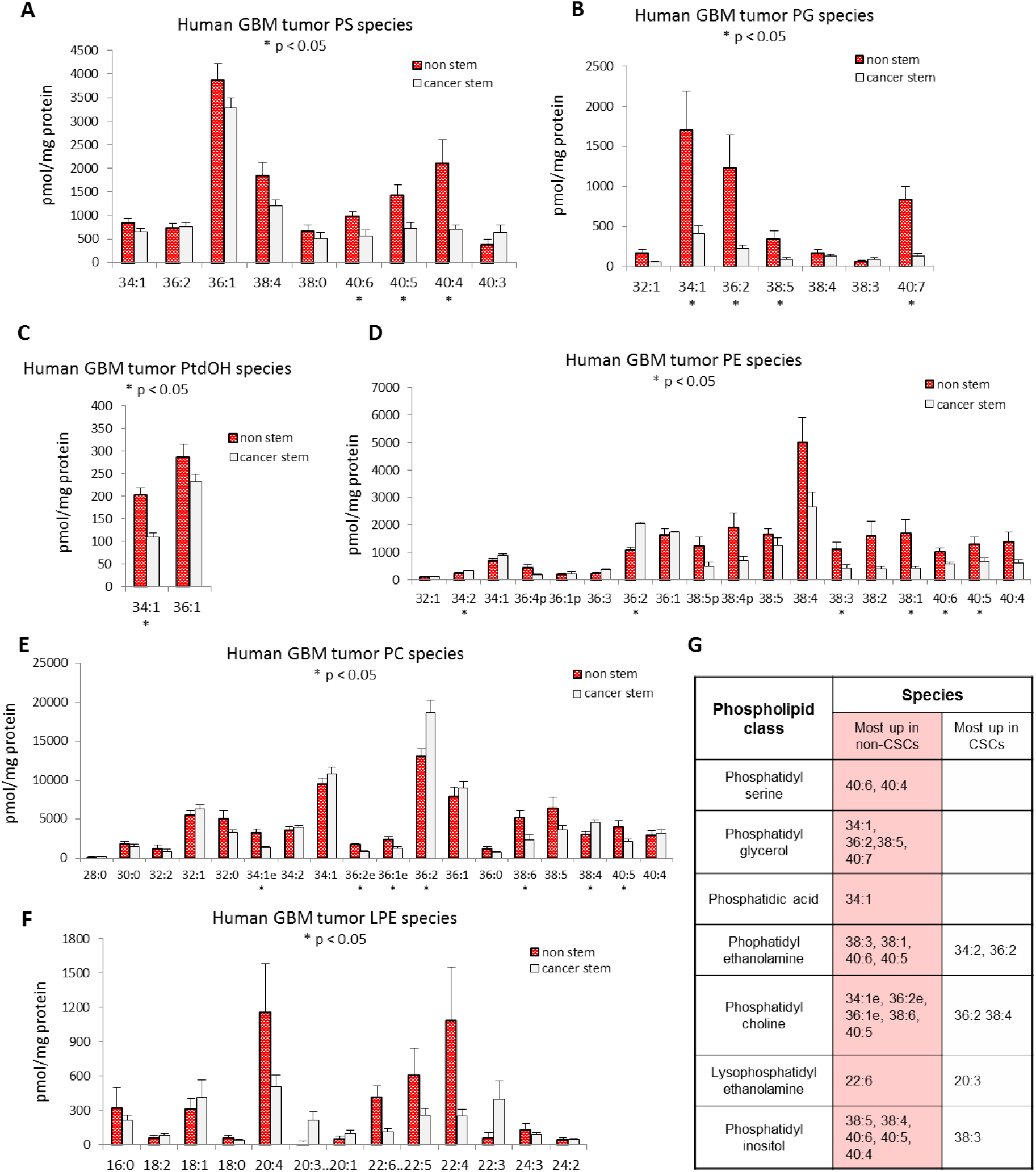
Lipidomic profiling of GBM stem and non-stem cells shows differences in specific lipid species. GBM CSCs and non-CSCs have differentially expressed phospholipid species, with most of the increase in lipid species in non-CSCs. (A) Phosphatidylserine, (B) Phosphatidylglycerol, (C) Phosphatidic acid, (D) Phosphatidylethanolamine, (E) Phosphatidylcholine, and (F) Lysophosphatidylethanolamine species. (G) Table highlighting the (statistically) significantly differences in lipid groups. * p < 0.05; *ns, p > 0.05*.

### CSCs maintain increased PUFA synthesis but not PUFA accumulation and storage through upregulation of fatty acid desaturase enzymes

Within the PI class, we observed a marked decrease in 38:4 PI and a reciprocal increase in 38:3 PI (Fig. 6A). It is generally accepted that 38:4 PI contains arachidonic acid (AA; 20:4; n-6) in the sn-2 position, while 38:3 PI is expected to harbor di-homo-γ-linolenic acid (DGLA; 20:3; n-6) in the sn-2 position. This reciprocal alteration prompted us to examine the expression levels of the delta-5 desaturase enzyme fatty acid desaturase 1 (FADS1), which converts DGLA to AA. Interestingly, the expression of FADS1 and that of the delta-6 desaturase FADS2 was elevated in CSC populations (Fig. 6B, C). We also found that both FADS1 and FADS2 were significantly upregulated in the organoid rim region and corresponding cellular tumor regions in patient tumors (Fig. 6D, E). A recent report showed that another PUFA-synthesizing enzyme, fatty acid elongase 2 (ELOVL2), exhibits a similar distribution^39^. Increased ELOVL2 and FADS2 further process AA into long-chain PUFAs, thus preventing high AA accumulation in CSCs despite high PUFA pathway activity. Given these findings, there is growing evidence for a role for PUFA synthesis in supporting tumorigenesis within the GBM microenvironment. The current evidence suggests that the generation of PUFA-enriched glycerophospholipids appears to be favored in the nutrient-rich CSC microenvironment and supported by high FADS2, FADS1, and ELOVL2 expression, whereas lipid accumulation and storage are favored in the nutrient-low non-CSC microenvironment (Fig. 6F). Taken together, our results demonstrate a striking degree of metabolic diversity in GBM depending on each cell’s microenvironment and CSC state, and this heterogeneity must be taken into account in both basic research and in therapeutic targeting of GBM.

**Figure 6.**
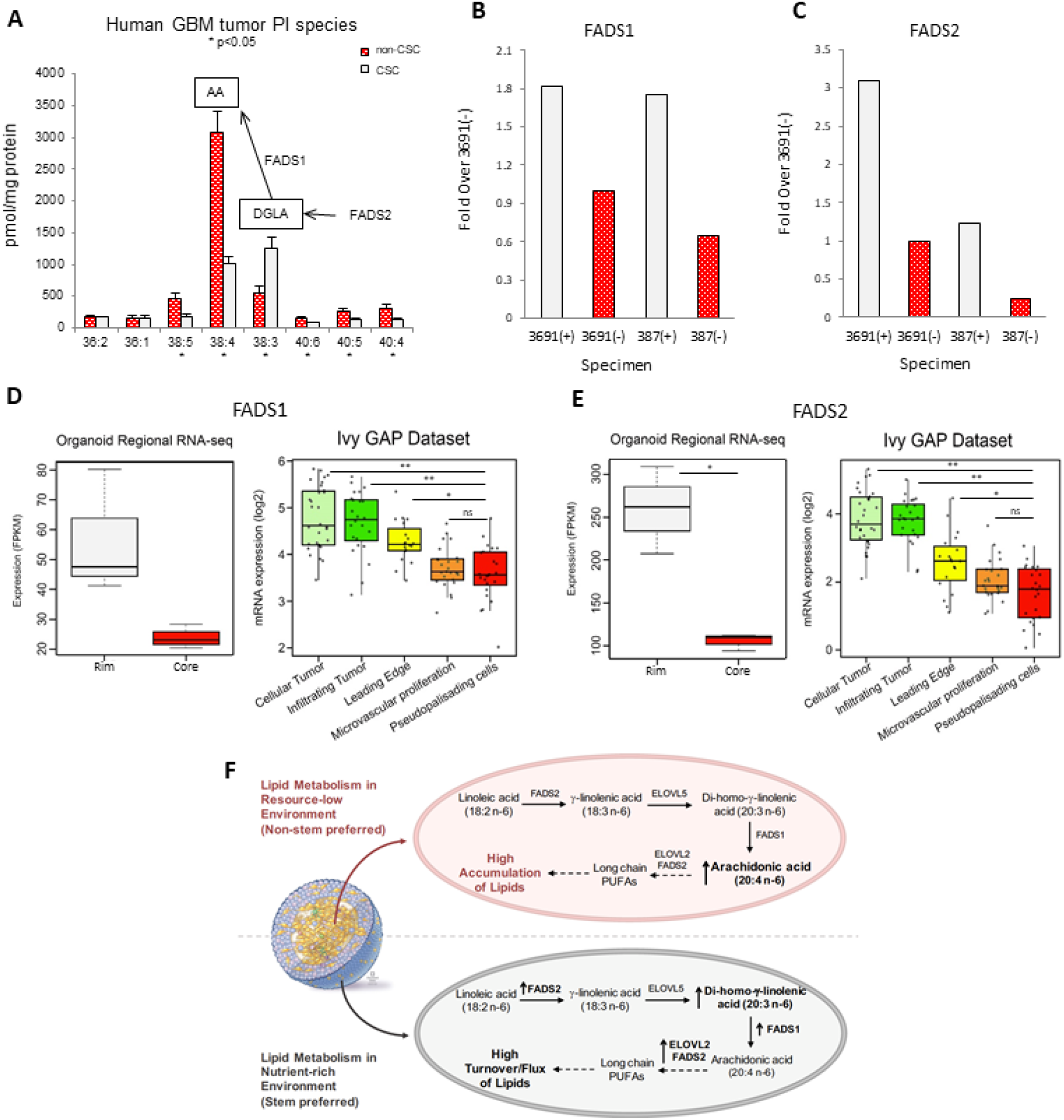
CSCs maintain increased PUFA synthesis but not PUFA accumulation and storage through upregulation of fatty acid desaturase enzymes. (A) GBM CSCs and non-CSCs have notable differences in phosphatidylinositol species. In particular, CSCs have reduced arachidonic acid (AA) levels but increased levels of its precursor DGLA. qRT-PCR shows that FADS1 and FADS2 levels are higher in CSCs compared to non-CSCs from PDX models (B) and (C). RNA-seq shows higher expression of FADS1 and FADS2 in cells of the GBM organoid proliferative rim and patient cellular tumor regions (D) and (E). (F) Proposed mechanism of high accumulation vs high flux of lipids in GBM non-CSCs and CSCs, respectively.

## DISCUSSION

To meet energy demands in resource-sparse tumor microenvironments, tumor cells undergo metabolic reprogramming, which is known as a hallmark of GBM in addition to many other cancers.^8,40^ Metabolic targeting has been proposed as a therapy for many tumor types, and inhibition of DGAT1 has recently been proposed to alter fat metabolism and increase oxidative stress in GBM.^11^ However, cellular heterogeneity and plasticity are features of GBM and drive therapeutic resistance.^41^ Recently, intratumoral heterogeneity has become a highly researched focus of both pre-clinical and clinical GBM studies aiming to develop targeted treatment methodologies^42^, and it is critically important to mimic this feature in in vitro cultures for more relevant study outcomes. Our overall findings show that lipid droplets accumulate in the hypoxic core of GBM organoids and also in perinecrotic and pseudopalisading regions of GBM patient tumors. This was accompanied by overall increased accumulation of fatty acid species in the CD133-negative non-CSC population versus matched CD133-positive CSCs obtained from patient-derived xenografts. In short, we show that intratumoral lipid metabolism heterogeneity exists and must be considered at the pathologic, cellular and molecular levels.

Pioneering studies by Patel et al.,^3^ and more recently by Neftel et. al,^36^ used single-cell RNA sequencing (scRNA-seq) to show that GBM cells vary in their expression of different transcriptional programs, including those driving proliferation and hypoxia. Consistent with these studies, we found that distinct meta-module signatures from patient tumor cells are enriched within our distinct GBM 3D organoid spatial regions. While cells from the organoid core highly express genes belonging to the mesenchymal-like (MES) meta-module, cells from the rim have higher expression of the astrocyte-like (AC) and oligodendrocyte-progenitor (OPC)-like meta-modules. Although spatial information is lost when tumors are dissociated for scRNA-seq, it is assumed that these different populations from patient tumors derive from different microenvironments within the tumor. Here, we further showed that the gene expression in organoids corresponds to the gene expression in the regional Ivy GAP database.^37^ The genes enriched in the organoid core reflect the patient pseudopalisading and perinecrotic tumor regions in the Ivy GAP data, whereas the genes enriched in the organoid rim were associated with the gene enriched in the cellular tumor region. Thus, we successfully ascertained that molecular heterogeneity in the 3D organoid culture corresponds to primary GBM tumors at both the single-cell and spatial microenvironmental levels.

We show that HILPDA expression, which is canonically driven by HIF-1α, is consistently increased in primary GBM tumors and specifically in the pseudopalisading/perinecrotic region of GBM (Fig. 2A). This supports a finding by Mao et al. showing that HILPDA was upregulated in GBM compared to normal brain tissue or lower-grade gliomas and in GBM cells cultured in hypoxic conditions.^43^ Additionally, studies show that HILPDA is involved in triglyceride fatty acid secretion^44^ and regulates lipid metabolism and hypoxia-induced lipid droplet biogenesis.^45^ When we investigated lipid droplet accumulation in our 3D GBM organoid model, we discovered that lipid droplets were exclusively enriched in the hypoxic core region of the 3D organoids. We further traced this finding back to patients, showing that all patients in our panel with clear pseudopalisading necrosis by pathology, and most patients overall, accumulate lipid droplets in the hypoxic regions of their tumors. To our knowledge, this is the first concrete demonstration of lipid droplet accumulation as a marker of cells surrounding pseudopalisading necrosis.

It is an outstanding open question in the field^8^ whether there is a link between self-renewal and lipid metabolism in GBM. Here, we addressed this question using GBM patient tissue, fresh patient-derived 3D ex vivo cultures, and patient-derived xenografts. We obtained consistent findings at the single molecular, transcriptional, cellular, and tissue scales, all of which show increased lipid content in nutrient-poor and non-stem GBM cells. Our results support findings showing that slower-cycling GBM cells (as found in GBM organoid cores (Fig. 1D and previous work^25^)) have increased lipid droplet content^46^ and that targeting lipid homeostasis in GBM has antiproliferative effects^11^. However this also contrasts with findings that slow-cycling tumor cells may be CSCs^46^ and that cultured colorectal cancer CSCs have increased lipid content compared to non-CSC populations^47^.

Along with lipogenesis, lipolysis (through oxidation of fatty acids) is critical for the renewal of stem cells and maintenance of stemness^48^, which could explain the decreased lipid accumulation and overall decrease of lipids in CSCs despite high lipid synthesis rates (Fig. 3D). One initial paradox in our data was the increased 20:4 AA species in non-CSCs despite lower DGLA (20:3) species compared to CSCs. Upon examining the expression levels of the fatty acid desaturase genes involved in AA/DGLA catabolism, *FADS1* and *FADS2*, we found that both are enriched in the CSC-rich organoid rim. Additionally, ELOVL2, a critical enzyme downstream of AA, has been shown to be enriched in SOX2- and OLIG2-positive GBM cells^39^. We propose that ELOVL2 and FADS2 in CSCs facilitate turnover/utilization of lipid species in high nutrient conditions, unlike in non-CSCs and GBM cells in nutrient-poor environments, where the fatty acids accumulate, forming lipid droplets. We believe that this potential mechanism makes intuitive sense from a standpoint of survival, where energy storage is favored in environments where nutrient resources are scarce and uncertain.

## Dedication

This work is dedicated in loving memory to the late Dr. H. Alex Brown (Vanderbilt School of Medicine), who passed away during the preparation of this manuscript. Dr. Brown’s work advanced the concept that alterations in lipid metabolism can support cancer initiation and progression.

## Acknowledgments

We thank Dr. Erin Mulkearns-Hubert for insightful discussion and constructive comments on the manuscript. We thank Mary McGraw and Pengjing Huang of the Rose Ella Burkhardt Brain Tumor Bank for their help with sample collection and clinical diagnostic information. We thank Amanda Mendelsohn (Cleveland Clinic for Medical Art and Photography) for illustration assistance. We thank the LRI flow cytometry core for FACS assistance.

