## Supplemental Figures 1-4 for "Altered lipid metabolism marks glioblastoma stem and non-stem cells in separate tumor niches"

**Supplemental Figure Captions**

**Figure S1. Marker gene expression of cells in the organoid core is comparative to the cells in the corresponding pseudopalisading region of primary patient GBM.** (A-D) Gene expression levels of potential regional or subtype marker genes across the TCGA, Gravendeel, Bao, and Ivy GAP datasets compared to GBM organoid regional sequencing data.

**Figure S2. Oil Red O staining of GBM organoids shows lipid droplet accumulation in the organoid core region of multiple independent GBM specimens.** Oil Red O staining of frozen sections of patient-derived GBM organoids at wide-field and high-power magnification (insets, representative for rim and core).

**Figure S3. Oil Red O staining in primary patient tumors shows enriched lipid droplet accumulation in pseudopalisading regions of multiple independent GBM specimens.** Oil Red O staining of frozen sections of patient-derived GBM organoids at wide-field and high-power magnification (insets, representative for cellular or pseudopalisading histology).

**Figure S4. Some phospholipid classes show equal abundance in CSCs compared to non-CSCs**. Quantitation of sphingomyelin (SM) lipid species from sorted CSCs and non-CSCs. No significant differences detected.

**Figure S1.** Marker gene expression of cells in the organoid core is comparative to the cells in the corresponding pseudopalisading region of primary patient GBM.

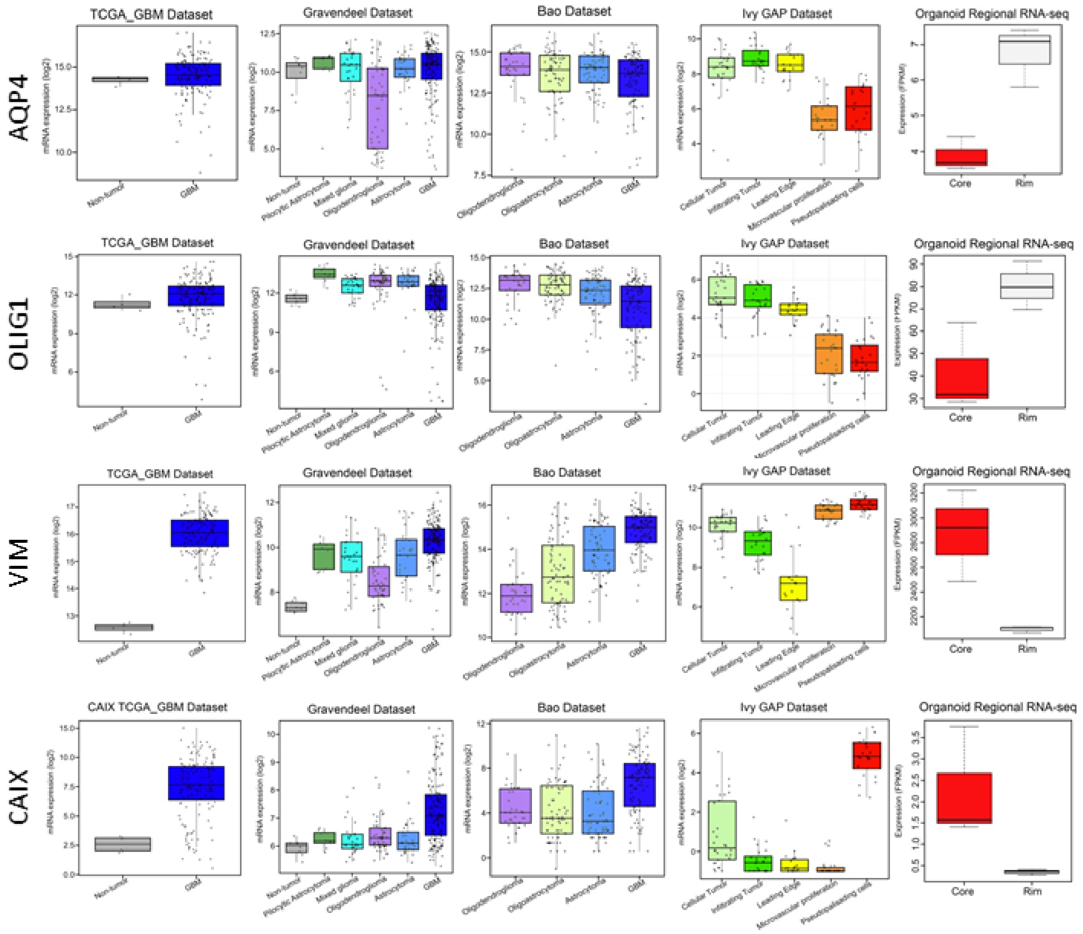

**Figure S2.** Oil Red O staining in 3D organoids shows enriched lipid droplet accumulation in organoid core regions.

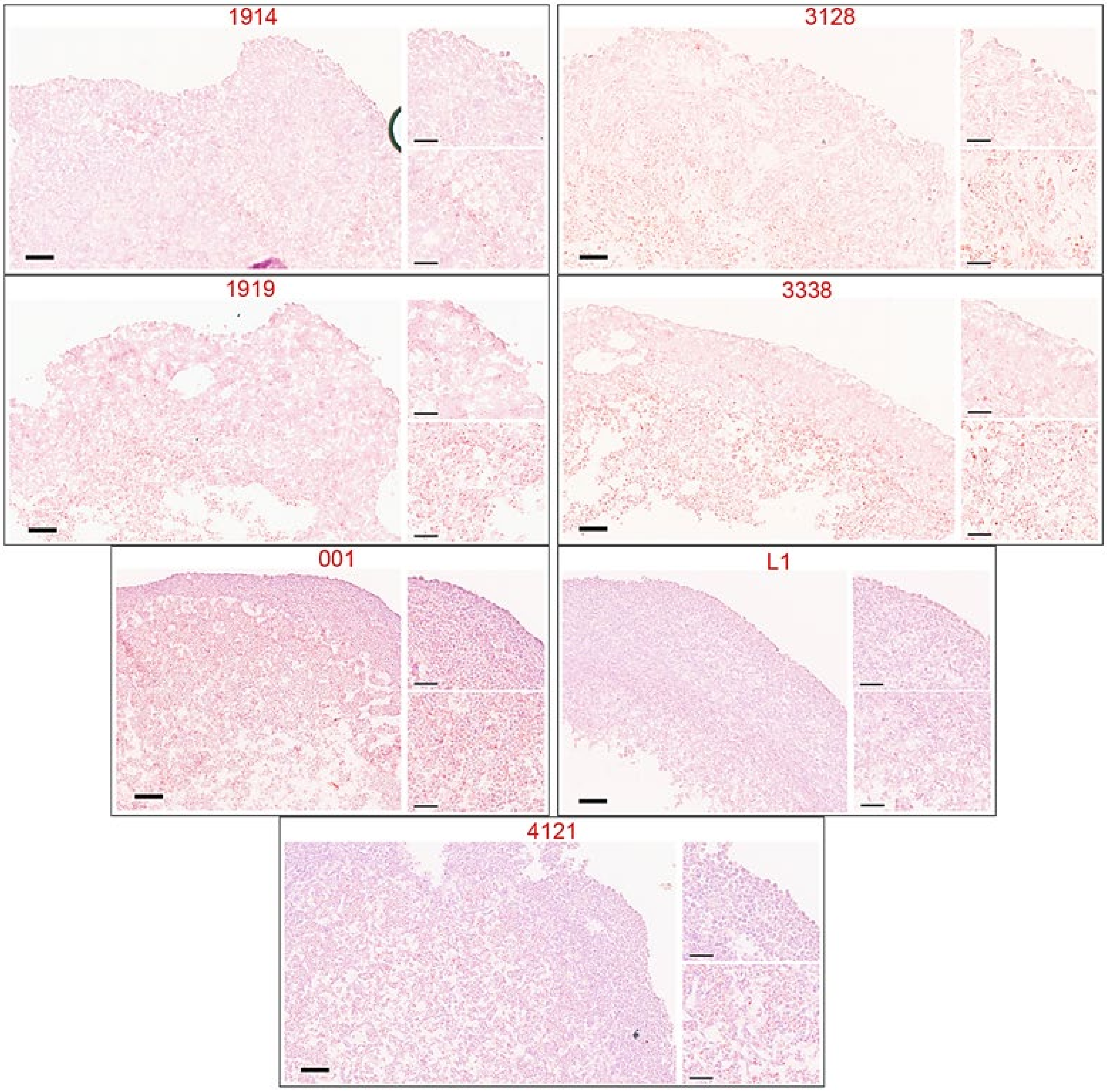

**Figure S3.** Oil Red O staining in primary patient tumors shows enriched lipid droplet accumulation in pseudopalisading regions.

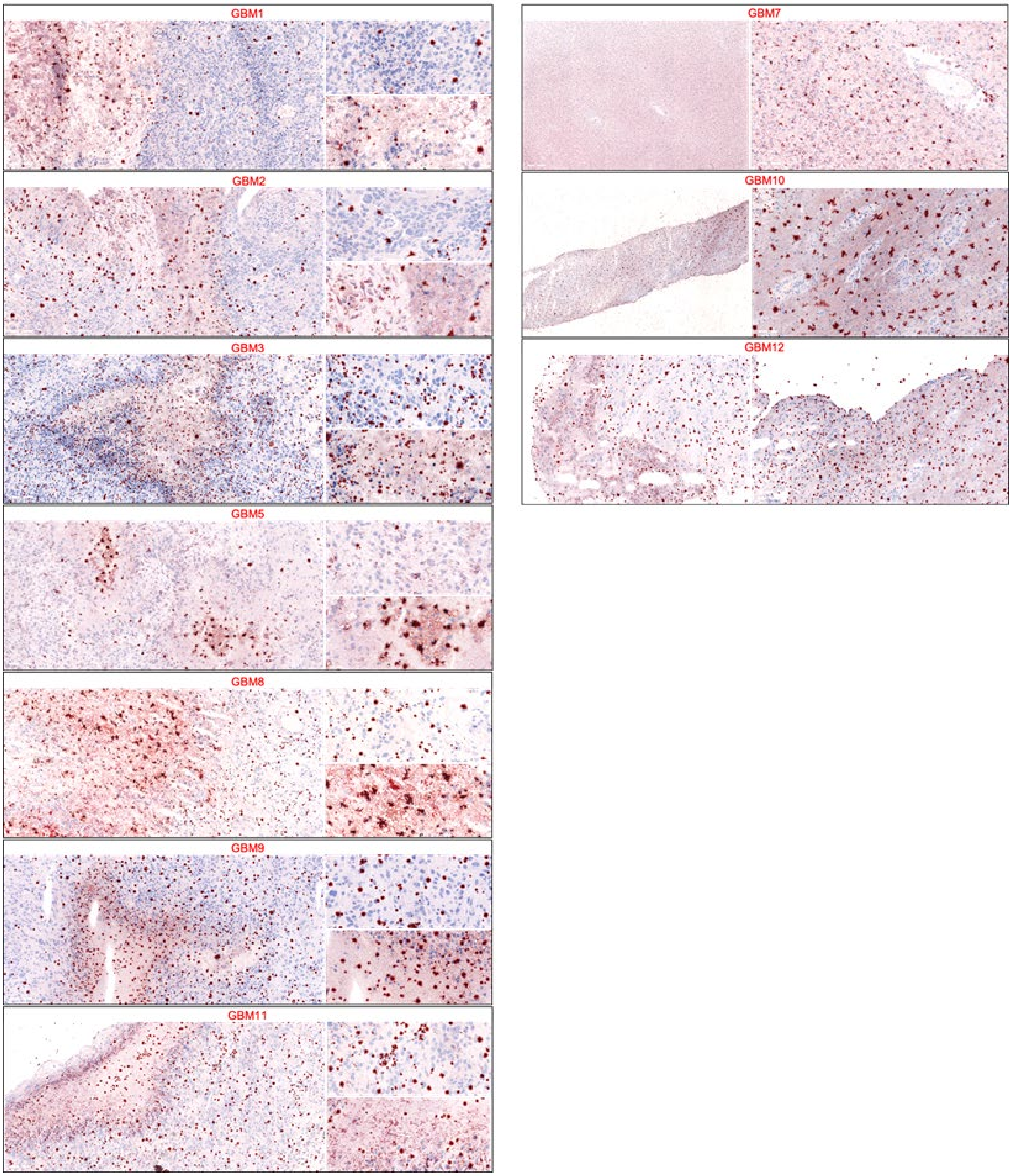

**Figure S4.** Some phospholipid classes show equal abundance in CSCs compared to non-CSCs.

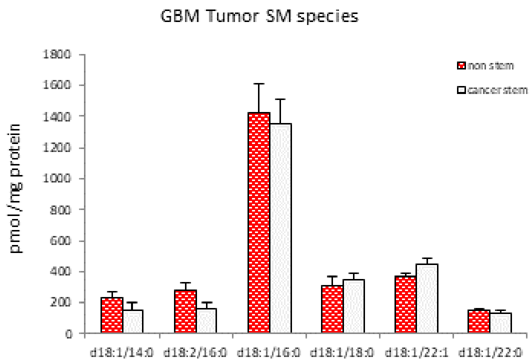
